# Re-assessing the phylogenetic status and evolutionary relationship of Forest Owlet [*Athene blewitti* (Hume 1873)] using genomic data

**DOI:** 10.1101/2021.10.09.463762

**Authors:** K.L Vinay, Meghana Natesh, Prachi Mehta, Rajah Jayapal, Shomita Mukherjee, V.V. Robin

## Abstract

Phylogenetic relationships are often challenging to resolve in recent/younger lineage when only a few loci are used. Ultra Conserved Elements (UCE) are highly conserved regions across taxa that help resolve shallow and deep divergences. We utilized UCEs harvested from whole genomes to assess the phylogenetic position and taxonomic affiliation of an endangered endemic owlet in the family Strigidae – the Forest Owlet *Athene blewitti*. The taxonomic placement of this species has been revised multiple times. A multigene study attempted to address the question but showed a discrepancy across datasets in its placement of the species within genus *Athene*. We assembled a dataset of 5018 nuclear UCE loci with increased taxon sampling. Forest Owlet was found to be an early split from the *Athene* clade but sister to other *Athene;* and consistent across three approaches - maximum likelihood, bayesian, and the multispecies coalescence. Divergence dating using fossil calibrations suggest that the *Athene* lineage split from its ancestor about 7.6Mya, and the Forest Owlet diverged about 5.2Mya, consistent with previous multigene approaches. Despite osteological differences from other *Athene*, we suggest the placement of the Forest Owlet as a member of the *Athene* to emphasize its evolutionary relationship.

**Graphical Abstract:** 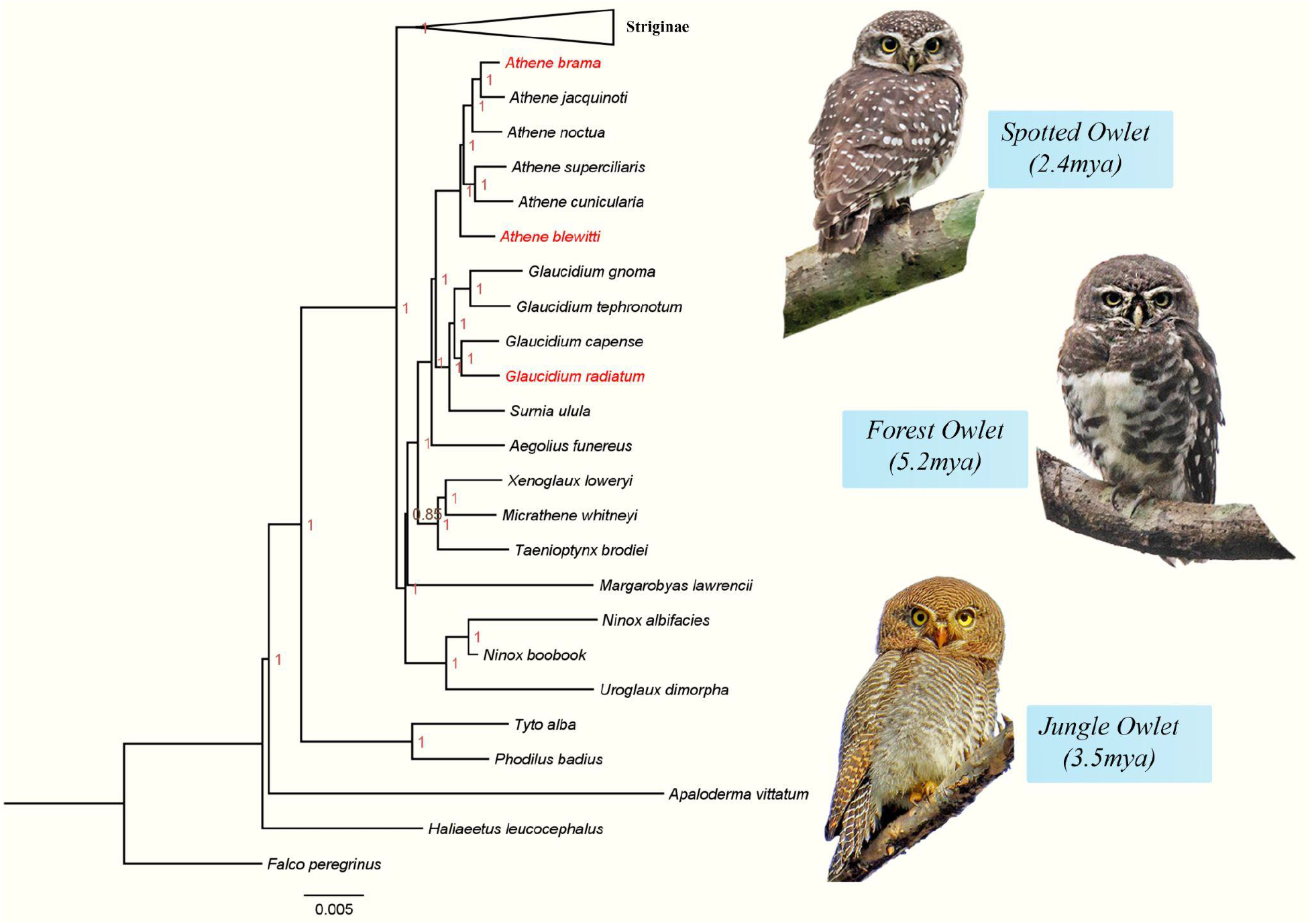

**HIGHLIGHTS:**

1. Phylogenomics using genome-wide nuclear markers yielded a well-supported topology for *Athene* and *Glaucidium* lineages.
2. Three different methods of phylogenetic tree construction showed that Forest Owlet is an early split from all other *Athene* species.
3. Divergence dating in the bayesian framework puts the Forest Owlet age between 5.0my to 5.5my.

## 1. INTRODUCTION

The hierarchy of systematic classification of organisms into taxa is based on the various biological pieces of evidence, i.e. morphology, genetics, palaeontology, etc. Taxonomic revisions to the existing classification arise with the addition of new species and understanding the relationships between the sister species (Kennedy et al., 2005). Information on biodiversity is vital for ecological and environmental assessments and subsequent evolutionary studies (Meredith et al., 2019). Mitochondrial markers and whole mitogenomes have been used extensively with nuclear markers to establish phylogenetic relationships among the species (Mueller, 2006). However, mitochondrial data have exhibited strong biases in phylogenetic placements across taxa (Luo et al., 2012; Zhong et al., 2011). In recent years, exomes (Wang et al., 2020) and Ultra Conserved Elements (UCE) have been used in phylogenetic analyses to resolve the nodes across taxa (Jarvis et al., 2014; Oliveros et al., 2019). The Tetrapod UCEs are conserved over millions of years across taxa, including birds (Faircloth et al., 2012), mammals (McCormack et al., 2012), and amphibians (Newman and Austin, 2016). These UCEs with variable flanking regions have proven helpful in resolving deep and shallow phylogenetic relationships within and across species (Erickson et al., 2020; Smith et al., 2014), including those that multigene phylogeny failed to resolve (Gilbert et al., 2015; McCormack et al., 2012). In some cases, the data type (Multigenes, Introns, UCE) has a more significant influence on the tree’s topology than the taxon sampling (Reddy et al., 2017).

The Forest Owlet is an endangered species endemic to central India, belonging to the typical owl family Strigidae and rediscovered in 1997 (King and Rasmussen, 1998). Its phylogenetic placement has always been under debate among taxonomists (Koparde et al., 2018). In 1873 A.O Hume described the species under genus *Heteroglaux* owing to the difference in the nostril morphology and smaller head compared to *Athene* (Hume, 1873). However, Ali and Ripley (Ali and Ripley, 1983) treated the Forest Owlet as *Athene*, as did a few others, including Monroe and Sibley (1997) and Roche (2000). In 2013 Forest Owlet was placed back in the genus *Heteroglaux* based on the collected specimens’ morphological and osteological features (Rasmussen and Collar, 2013). Later in 2018, a multigene study using two mitochondrial genes and three nuclear genes suggested that Forest Owlet is nested within the *Athene* clade (Koparde et al., 2018). This was also followed in the International Ornithological Committee bird list (IOC World Bird List v8.2, 2018) and Birds of the World (Holt et al., 2020). However, there was a discordance between mitochondrial and nuclear genes with regard to the placement of Forest Owlet within the genus *Athene* (Koparde et al., 2018). Hence, in this study, we proposed to investigate the Forest Owlet’s phylogenetic placement, evolutionary relationships and re-assess relationships with other *Athene* species with additional data, using several thousand Ultra Conserved Elements.

## 2. OBJECTIVES

Assess the phylogenetic status of the Forest Owlet *Athene blewitti* with genomic (Ultra Conserved Elements -UCE) data and determine its date of divergence.

## 3. METHODS

### 3.1 DNA extraction and sequencing

DNA was extracted from blood collected from adult Forest Owlet (with due permits) with Qiagen DNeasy Blood and Tissue kit following the manufacturer’s protocol, with minor alterations. The extract was quantified using a Qubit 4 fluorometer and screened on a 1% agarose gel. DNA was sequenced on the Illumina NovaSeq 6000 sequencer (150bp paired-end reads), targeting 100X coverage.

### 3.2 Analyses

#### Harvesting UCE and collating existing UCE data

Raw sequence reads were inspected for overall quality, adapter content, and the number of reads to ensure they met the required quality standards (TrimGalore!^1^). Reads were filtered to remove poor-quality or uncalled bases and adapters before assembling into contigs *denovo* using Megahit 1.2.9 (Li et al., 2016). Ultra Conserved Elements loci were harvested following Phyluce tutorial III (Faircloth, 2016). Assembled contigs of previously sequenced UCE data for the Strigidae family and outgroups were obtained from Dryad (Salter et al., 2020). Spotted Owlet and Jungle Owlet whole-genome sequences generated for another study (Natesh et al., 2020) were also denovo assembled and used to harvest the UCEs, as described above. Next, we followed the Phyluce Tutorial I (Faircloth, 2016) to align, trim and concatenate the data for the downstream process. For this study, we renamed taxa following the IOC World Bird list taxonomy (IOC World Bird List v11.2, 2021). Detailed methods are explained in the supplementary materials.

### 3.3 Phylogenetic analyses with UCE data

The concatenated sequences were exported to phylip and nexus formats for further analyses. We used three approaches to reconstruct trees, as described below. For the Maximum Likelihood and Bayesian trees, we used the GTR GAMMA model. For the Maximum Likelihood tree (RAxML 8.2; Stamatakis, 2014), 20 searches were done for the best tree, which was then combined with 1000 bootstrap replicates. For the Bayesian tree, we used ExaBayes 1.5 (Aberer et al., 2014) to perform four independent runs of 2 Markov chains for 2 million generations each. In addition, we also used a multispecies coalescent approach - SVDquartets, implemented in PAUP* 4.0a16 (Swofford and Sullivan, 2003). This method uses only single site-based information and helps side-step issues related to incomplete lineage sorting and estimation of gene trees (Chifman and Kubatko, 2014). Details of the runs and model convergence are described in the supplementary materials. All the trees were visualised in FigTree v1.4.3 (Rambaut, 2012).

### 3.4 Divergence time dating

We estimated divergence dates in a Bayesian framework using the MCMCTree program (PAML 4.8; dos Reis and Yang, 2011). This uses the Taylor expansion to avoid calculating likelihood at every MCMC iteration for the divergence time estimation (dos Reis and Yang, 2011). We used the gene tree from Bayesian inference as the input gene tree. Two priors, the overall substitution rate (*rgene gamma*) and rate drift parameter (*sigma2 gamma*), were set to G (1 58.2) and G (1 1.2), respectively, with independent rates clock model (Rannala and Yang, 2007). The Strigidae family has several fossil records, especially from North America and Europe (Kurochkin and Dyke, 2011). We included three fossil records which are spread across the clades/tribes, i.e. oldest *Athene* fossil record (3.6My-5.4My) (Pavia et al., 2014), MRCA of Strigiformes (63.5-68.5My) (Kurochkin and Dyke, 2011) and the common ancestor of *Bubo scandiacus* and *Bubo nipalensis* (4my) (*König et al., 2008*) It is argued that using multiple fossil constraints with upper and lower bounds gives a more rational divergence estimation time (Benton and Donoghue, 2007; Inoue et al., 2010). We used the *mcmctree_tree_prep* package^2^ to include fossil constraints in the tree and used it as input for MCMCtree. Tracer 1.7 (Rambaut et al., 2018) was used to check for the convergence of the likelihood and other parameters. The output was visualised in FigTree v1.4.3 (Rambaut, 2012). Detailed methods and settings are described in the Supplementary Materials.

## 4. RESULTS

Sequencing yielded 614.5 million raw reads for the Forest Owlet. UCE harvesting from the denovo assembled contigs resulted in 4920, 4953, and 4948 loci for Forest Owlet, Jungle Owlet and Spotted Owlet, respectively (Detailed statistics for denovo assembly (Table1) and UCEs (Table2) in Supplementary materials). Across taxa, we enriched 5018 loci shared by at least 38 taxa, which resulted from 4326 alignments. The 75% complete concatenated matrix had 2,240,927 bp and 121,747 informative sites.

Concatenated data (ML and Bayesian -Supplementary Figure 1 & 2) and Multispecies Coalescent (SVD Quartets - Supplementary Figure 3) based methods produced identical tree topologies, and the relations among the species remained the same for the *Athene* and *Glaucidium* lineages, with high node supports (Figure 1). *Athene brama* and *Athene jacquinoti* share a common ancestor and are sisters to *Athene noctua*, whereas *Athene superciliaris* and *Athene cunicularia* are sister species. Using thousands of conserved nuclear loci in all three tree construction methods, Forest Owlet was recovered as an early split from all other *Athene* species. Our divergence dating analysis (Figure 1) recovered a 5.25my divergence of *Athene blewitti* from its most recent ancestor (95% HPD 5.0my-5.5my), whereas the *Athene brama* and *Athene jacquinoti* diverged about 2.4mya (95% HPD 1.3my-3.6my). *Athene noctua*, a sister to *Athene brama* and *Athene jacquinoti*, is 3.6my old (95%HPD 2.5my-4.6my). *Athene cunicularia* and *Athene superciliaris* split about 3.6mya (95% HPD 2.2my-4.5my).

**Figure 1:**
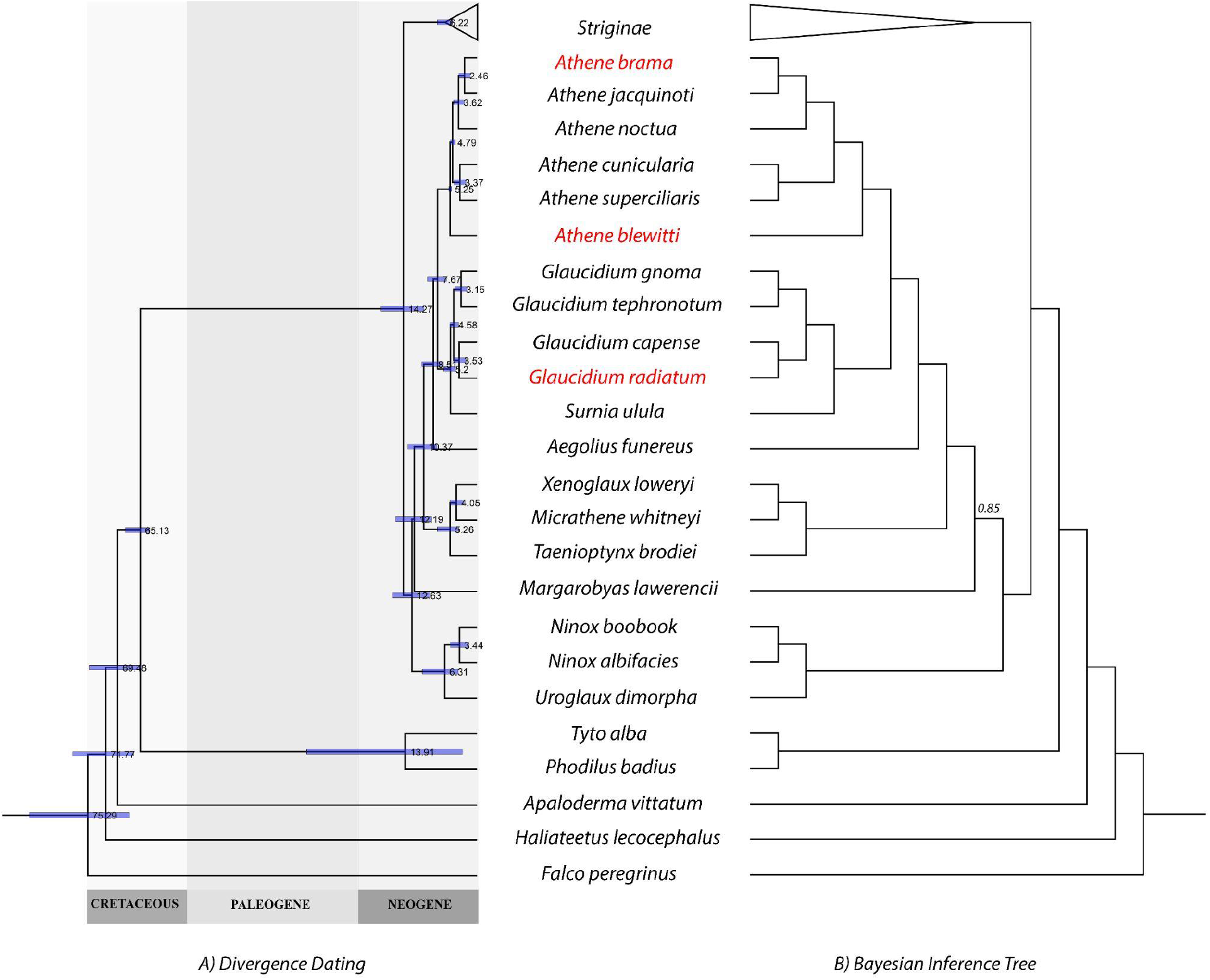
Phylogram of the divergence dating (A) and Cladogram of Bayesian Inference (B). All nodes have 100% posterior probability if not indicated otherwise. Node bars represent 95%HPD. Species in red colour are the ones included in this study. Collapsed species broadly belong to the Striginae clade.

*Glaucidium radiatum* and *Glaucidium capense* split about 3.5mya (95% HPD 2.2my-4.6my), and the entire Strigidae family diversified about 65.1Mya. (95% HPD 63.3my-68my).

## 5. DISCUSSION

In this study, we reconstruct maximum-likelihood, Bayesian inference, and coalescence-based phylogenies to infer the phylogenetic position of the Forest Owlet, with respect to the Athene clade, within the Strigidae group. Our analysis contradicts some findings of previous phylogenetic assessments but reaffirms others. We find that the Forest Owlet is an early split from the *Athene* clade, whereas, in Koparde et al., 2018, it is nested within *Athene*, either as sister to the clade from Madagascar Americas or to the Eurasian *Athene* species. We recovered the Spotted Owlet *Athene brama* as sister to Solomon’s Boobook *Athene jaqcuinoti* in all tree construction methods. Two previous studies, (Koparde et al., 2018; Sun et al., 2020), suggested that it is sister to the Little Owl *Athene noctua* since *A. jacquinoti* was not included in that study. Nevertheless, our divergence date estimates using UCE data agree with the previous estimates for the *Athene* group and Strigidae family (Jarvis et al., 2014; Koparde et al., 2018; Prum et al., 2015). This study adds considerable data over previous assessments and adds two *Athene* species to the global phylogeny. Therefore, it is unlikely that the topology for the *Athene* clade will change in subsequent analyses without substantially different methods.

However, the genus *Glaucidium* (Pygmy Owls), regardless of the inclusion of Jungle Owlet, remains under-represented (15% complete) and requires further evaluation.

Species with small ranges and specific niche requirements may have unique evolutionary histories, and Forest Owlet is one such species (Koparde et al., 2018) Estimating robust phylogenetic relationships is thus central to understanding evolutionary relationships (Edwards, 2009). However, ecological and morphological characters are also highly informative and can provide substantial insights. The Forest Owlet shares traits with other species in the genus *Athene* but has differences as well. The extreme similarities in the plumage of Forest Owlet with the Spotted Owlet, an *Athene*, could be a case of *symplesiomorphy*. The *Athene* clade, as circumscribed here sensu lato, is also marked by wide variations in activity time, ranging from primarily nocturnal (*Athene superciliaris*), and largely crepuscular (*Athene brama, Athene noctua*) to largely diurnal (*Athene cunicularia*) (König et al., 2008). Forest Owlet is known to be primarily diurnal and crepuscular. Osteological features and external morphological features such as heavily feathered toes, less spotted crown set the Forest Owlet apart from other *Athene* species, elucidated in detail by (Rasmussen and Collar, 2013), who chose to assign the species to *Heteroglaux*. Thus, weighing these aspects, taxonomically, the Forest Owlet can either be classified as a member of *Athene* or placed in its own monotypic genus *Heteroglaux*. However, we assign the species to *Athene*, as a resurrection of *Hetereoglaux* would render the remaining taxa under *Athene* paraphyletic unless *A. cunicularia and A. superciliaris* are also moved to monotypic genera.

Well-resolved phylogenies are the backbone for many ecological or evolutionary studies, including exploration of diversity, biogeography, evolutionary history, and adaptation. Among modern vertebrates, birds are one of the most diverse groups of organisms (Brusatte et al., 2015) and the quest to understand the evolutionary relationships among species has been ongoing. However, despite birds being a generally well-sampled group, South-Asian diversity remains under-represented in global phylogenies (Reddy, 2014), which may limit or even bias inferences from such datasets. UCE data are increasingly being used for greater resolution and provide comparable datasets across taxonomic groups.

## Acknowledgements

We are grateful to the State Forest Departments of Madhya Pradesh for granting us permission to collect samples (Permit Letter number: क्रमांक/मा.ची.-II/रि सर्च Date: 28.01.2021), the Scientific Computing Facility at IISER Tirupati and members of the IT Department for HPC access and help with troubleshooting. We thank Brant C Faircloth for helping us with the bioinformatics pipeline. We are also grateful to Frank E. Rheindt, Michael Wink and Per Alström for discussions, and their suggestions on the taxonomic placement of the Forest Owlet. We thank John McCormack and Whitney Tsai for helping us with the UCE lab pipeline. We thank Ashwin Warudkar, Naman Goyal and Pankaj Koparde for discussion and suggestions throughout this study. The Director, along with the Library, Administrative and Finance staff of SACON for logistic support. We thank Shashank Dalvi for providing us with the photos of Forest Owlet and Spotted Owlet, and we credit Shantanu Kuveskar (Shantanu Kuveskar, CC BY-SA 4.0 via Wikimedia Commons) for Jungle Owlet photograph to be used for graphical abstract.

## Funding

This study was funded from MoEFCC grant to Shomita Mukherjee, Rajah Jayapal and VV Robin (File Number: J.22012/61/2009-CS(W), Dated: 29th September 2017), and DST-SERB Early Career Grant to VV Robin (ECR/2016/001065)

## Data availability statement

De-novo assemblies, harvested UCEs, alignments and scripts used for the study are available from Open Science Framework (OSF) (Vinay et al, 2021).

## CRediT authorship contribution statement

VKL: Investigation, Data Curation, Formal Analysis, Visualization, Validation, Writing-Original draft, Writing-Review & Editing. MN: Supervision, Validation, Writing-Original draft, Writing-Review & Editing. PM: Resource, Writing-Review & Editing. RJ: Conceptualization, Writing-Review & Editing. SM: Conceptualization, Resource, Writing-Review & Editing, Supervision, Funding Acquisition. VVR: Conceptualization, Resource, Writing-Original Draft, Writing-Review & Editing, Supervision, Funding Acquisition.

## Declaration of Competing Interest

The authors declare that they have no known competing financial interests or personal relationships that could have appeared to influence the work reported in this study.

## Footnotes

1 https://www.bioinformatics.babraham.ac.uk/projects/trim_galore/

2 https://github.com/kfuku52/mcmctree_tree_prep

## Notes

https://github.com/stachyris/owlet_phylogeny

### Competing Interest Statement

The authors have declared no competing interest.

